# Tryptophan depletion sensitizes the AHR pathway by increasing AHR expression and GCN2/LAT1-mediated kynurenine uptake, and potentiates induction of regulatory T lymphocytes

**DOI:** 10.1101/2023.01.16.524177

**Authors:** Marie Solvay, Pauline Pfänder, Simon Klaessens, Luc Pilotte, Vincent Stroobant, Juliette Lamy, Stefan Naulaerts, Quentin Spillier, Raphaël Frédérick, Etienne De Plaen, Christine Sers, Christiane A. Opitz, Benoit J. Van den Eynde, Jing-Jing Zhu

## Abstract

**Background:** Indoleamine 2,3-dioxygenase 1 (IDO1) and tryptophan-dioxygenase (TDO) are enzymes catabolizing the essential amino acid tryptophan into kynurenine. Expression of these enzymes is frequently observed in advanced-stage cancers and is associated with poor disease prognosis and immune suppression. Mechanistically, the respective roles of tryptophan shortage and kynurenine production in suppressing immunity remain unclear. Kynurenine was proposed as an endogenous ligand for the aryl hydrocarbon receptor (AHR), which can regulate inflammation and immunity. However, controversy remains regarding the role of AHR in IDO1/TDO-mediated immune suppression, as well as the involvement of kynurenine. In this study, we aimed to clarify the link between IDO1/TDO expression, AHR pathway activation and immune suppression.

**Methods:** AHR expression and activation was analyzed by qRT-PCR and western blot analysis in cells engineered to express IDO1/TDO, or cultured in medium mimicking tryptophan catabolism by IDO1/TDO. *In vitro* differentiation of naïve CD4^+^ T cells into regulatory T cells (Tregs) was compared in T cells isolated from mice bearing different *Ahr* alleles or a knockout of *Ahr*, and cultured in medium with or without tryptophan and kynurenine.

**Results:** We confirmed that IDO1/TDO expression activated AHR in HEK-293-E cells, as measured by the induction of AHR target genes. Unexpectedly, AHR was also overexpressed upon IDO1/TDO expression. AHR overexpression did not depend on kynurenine but was triggered by tryptophan deprivation. Multiple human tumor cell lines overexpressed AHR upon tryptophan deprivation. AHR overexpression was not dependent on GCN2, and strongly sensitized the AHR pathway. As a result, kynurenine and other tryptophan catabolites, which are weak AHR agonists in normal conditions, strongly induced AHR target genes in tryptophan-depleted conditions. Tryptophan depletion also increased kynurenine uptake by increasing SLC7A5 (LAT1) expression in a GCN2-dependent manner. Tryptophan deprivation potentiated Treg differentiation from naïve CD4^+^ T cells isolated from mice bearing an AHR allele of weak affinity similar to the human AHR.

**Conclusions:** Tryptophan deprivation sensitizes the AHR pathway by inducing AHR overexpression and increasing cellular kynurenine uptake. As a result, tryptophan catabolites such as kynurenine, more potently activate AHR, and Treg differentiation is promoted. Our results propose a molecular explanation for the combined roles of tryptophan deprivation and kynurenine production in mediating IDO1/TDO-induced immune suppression.

**SIGNIFICANCE:** In preclinical models, tryptophan degradation by IDO1 or TDO was shown to induce tumoral resistance to immune rejection, by restricting inflammation and promoting T-cell tolerance to immunogenic tumor antigens. However, the mechanism that translates these metabolic changes into T-lymphocyte malfunction within the tumor microenvironment (TME) is still uncertain. It has been proposed that kynurenine, the main tryptophan catabolite, acts as an endogenous ligand for the aryl hydrocarbon receptor (AHR), leading to the suggestion that the IDO1/Kyn/AHR axis could play a key role in modulating inflammatory and immune responses. However, recent studies challenged the notion that kynurenine is a genuine and potent AHR agonistic ligand. Moreover, the relative role of tryptophan depletion versus kynurenine production in IDO1/TDO mediated immune suppression remains unknown.

In this work, we further explored and clarified the association between IDO1/TDO activity and AHR activation. Unexpectedly, we observed that tryptophan depletion strongly increased AHR expression, thereby potentiating its activation by weak agonists such as kynurenine and derivatives. Tryptophan depletion thereby potentiated the induction of regulatory T cells. This was particularly true in mouse strains that express an Ahr allele of weak affinity, similar to the human AHR.

Tryptophan depletion also increased cellular kynurenine uptake by increasing SLC7A5 (LAT1) expression in a GCN2-dependent manner, thereby also contributing to a better AHR activation by kynurenine upon tryptophan depletion.

Altogether, our findings identify a new mechanism explaining IDO/TDO mediated AHR activation and immune suppression, based on the sensitization of the AHR pathway by tryptophan depletion, resulting in a higher AHR stimulation by weak agonists of the kynurenine pathway, and a better induction of regulatory T cells.

## INTRODUCTION

Indoleamine 2,3-dioxygenase 1 (IDO1) and tryptophan-dioxygenase (TDO) are two enzymes responsible for catabolizing the essential amino acid tryptophan into kynurenine. Expression of these enzymes is frequently observed in advance-stage cancers and is associated with poor disease prognosis (1). In preclinical *in vivo* models IDO1 and TDO were found to induce tumoral resistance to immune rejection, by restricting inflammation and promoting T-cell tolerance to immunogenic tumor antigens (2, 3). Despite their ability to promote immune rejection of tumors in preclinical models, IDO1 inhibitors so far have failed to improve the outcome of immunotherapy in human melanomas (4). This calls for a better understanding of IDO1-mediated immune suppression in both mouse and human systems. Indeed, the mechanism that translates tryptophan catabolism into T-lymphocyte malfunction within the tumor microenvironment (TME) remains uncertain. Several hypotheses have been put forward to explain IDO1-mediated immune suppression, but today it remains unclear whether the major driver for suppression is the local depletion of tryptophan or the production of kynurenine and derivatives. One of the first models proposed was based on the observed proliferation arrest of T lymphocytes exposed to low (< 1 µM) tryptophan concentrations (5). This arrest was found to depend on an integrated stress response triggered by general control non-derepressible 2 (GCN2), a kinase that is sensitive to transfer RNAs that are not bound to their amino-acid cargo, whose levels increase upon amino-acid depletion (6). However, subsequent work showed that GCN2 was not involved in the IDO1-mediated tumoral immune resistance (7). Another model proposed that IDO1-mediated suppression relied on deactivation of the mTOR pathway due to tryptophan depletion (8). The alternative view that the production of kynurenine and derivatives is the major driver of immunosuppression was supported by initial observations that kynurenine and its catabolites could trigger apoptosis of T lymphocytes (9). Moreover, kynurenine itself was then identified as an endogenous ligand for the aryl hydrocarbon receptor (AHR) (10-12), leading to the suggestion that the IDO1/Kyn/AHR axis could play a key role in modulating inflammatory and immune responses (12, 13). Present in cells of various organs, AHR acts as an intracellular receptor sensitive to a variety of aromatic molecules potentially toxic to cells. Once activated, AHR is translocated into the nucleus and induces transcription of detoxifying enzymes, such as cytochrome P450 encoded by *CYP1A1*, involved in the catabolism of aromatic compounds (14). AHR activity promotes the induction of peripheral regulatory T lymphocytes (Tregs) (10, 15), the upregulation of PD-1 on CD8^+^ T cells (16) and the recruitment of immunosuppressive tumor-associated macrophages (TAM) (17). However, despite considerable advances in the understanding of immunosuppressive mechanisms mediated by AHR, a recent study challenged the notion that kynurenine is a genuine AHR agonistic ligand. Seok et al. (18) observed that kynurenine dissolved in DMSO spontaneously undergoes chemical conversion into an oxidized dimeric derivative that is an exceptionally strong AHR agonist and whose trace presence in kynurenine preparations may account for the reported weak agonistic activity of kynurenine. This phenomenon was usually only observed at high concentrations well above the physiological concentration (18). In addition, the chemical structure of kynurenine does not make it an optimal ligand, as it is highly polar and also too small for good binding in the AHR ligand binding site, which rather accommodates ligands with more than one aromatic ring (18). Kynurenine metabolites such as anthranilic acid (AA), kynurenic acid (KA) and 3-hydroxykynurenine (HK), which have a structure more apt to bind AHR, have also been proposed as AHR ligands, but their concentration in human serum is 10-50 times lower than kynurenine (19).

In this work, we further explored and clarified the association between IDO1/TDO activity and AHR activation. Unexpectedly, we observed that tryptophan depletion strongly increased AHR expression, thereby potentiating its activation by weak agonists and the induction of regulatory T cells. These results provide a mechanistic rationale supporting a combined effect of both tryptophan depletion and production of tryptophan catabolites in the local immunosuppression triggered by IDO1/TDO activity.

## RESULTS

### IDO1 and TDO induce AHR activation and overexpression

We first aimed to confirm the correlation between the enzymatic activity of IDO1/TDO and the activation of AHR in human cells. As shown in Figure 1A and 1B, human embryonic kidney cells (HEK293-E) overexpressing IDO1 or TDO efficiently converted tryptophan into kynurenine (Fig 1A, B, right panels), and triggered AHR activity, as indicated by the upregulated expression of AHR-target genes *CYP1A1* (Fig 1A and 1B, left panels), *CYP1B1, STC2* (23), *TIPARP, EREG* and *IL4I1* (24) (Suppl Fig 1A). This AHR activation was dependent on IDO1/TDO enzymatic activity, as it was abolished in the presence of IDO1 or TDO inhibitors. Unexpectedly, in addition to AHR activation, we observed a strong increase in the expression of the *AHR* transcript itself, which was also dependent on the enzymatic activity of IDO1 or TDO (Fig 1A and 1B, middle panels).

**Figure 1.**
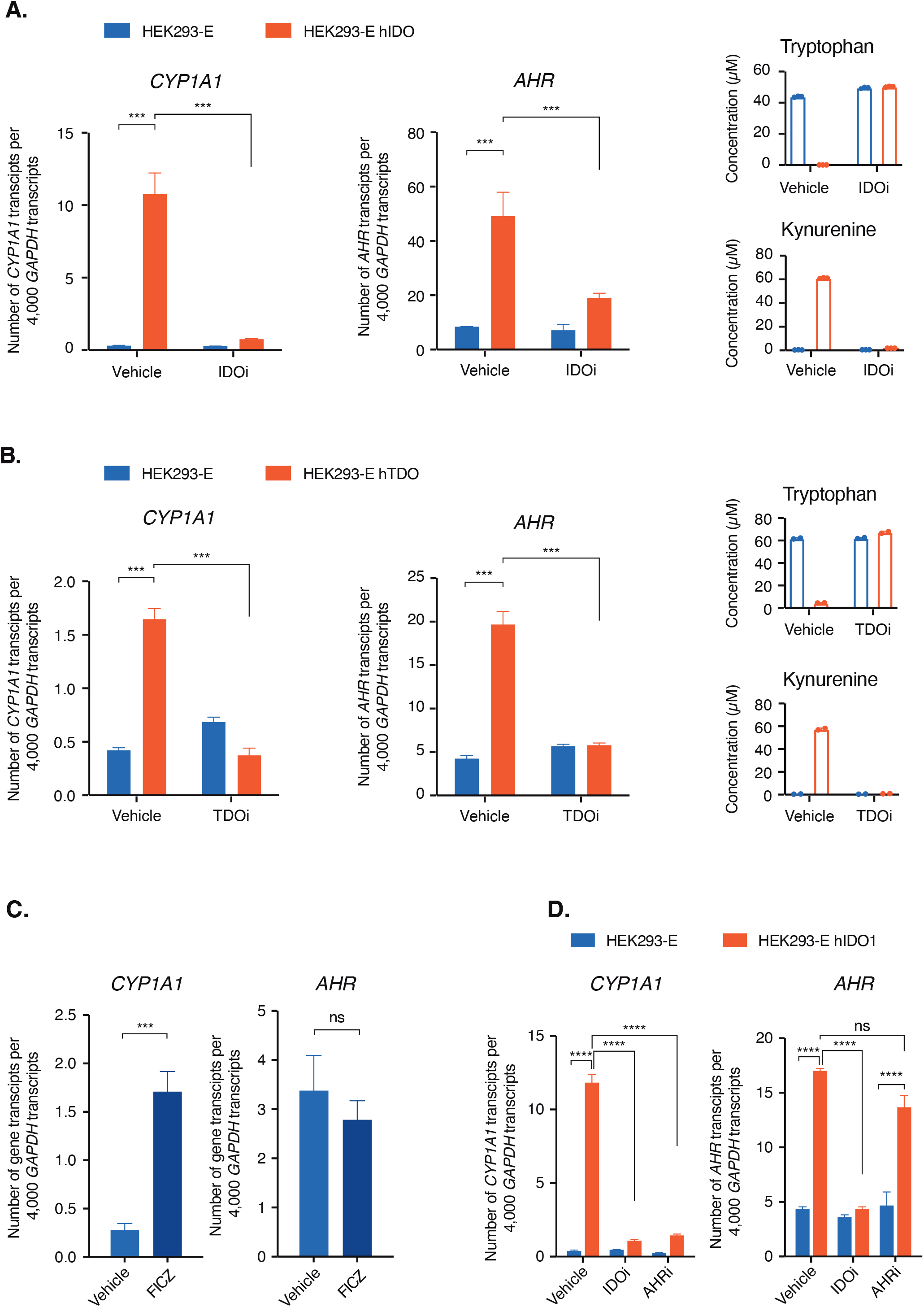
IDO1 and TDO activity induces AHR activation and upregulation. qRT-PCR analysis of *CYP1A1* (left panel) and *AHR* (middle panel) in HEK293-E or HEK293-E hIDO1 cells treated or not with IDO1 inhibitor (Epacadostat, 2.5 µM) for 120 h. qRT-PCR analysis of *CYP1A1* (left panel) and *AHR* (middle panel) in HEK293-E or HEK293-E hTDO cells treated or not with TDO inhibitor (EOS200809, 5 µM). Kynurenine or tryptophan concentrations were measured by HPLC in cell supernatant (A and B, right panels). **C**. qRT-PCR analysis of *CYP1A1* and *AHR* in HEK293-E cells treated with the AHR agonist FICZ (1 μM) for 120 h. **D**. qRT-PCR analysis of *CYP1A1* and *AHR* in HEK293-E or HEK293-E-IDO1 cells treated or not with IDO1 inhibitor (Epacadostat, 2.5 µM) or AHR inhibitor (CH-223191, 10 µM) for 120 h. The mRNA levels of different genes were measured by quantitative qRT-PCR and normalized to *GAPDH*. Mean ± SD of technical triplicates from one representative experiment out of three independent experiments. **** significant with p<0.0001, *** significant with p<0.001, ** p<0.01, * p<0.05 and NS= not significant.

Because a previous study suggested a possible “autoregulation” of AHR expression by AHR agonists in a positive feedback manner (25), we evaluated whether the increased AHR expression in IDO1/TDO expressing cells was dependent on AHR activity. However, we observed no increased AHR expression in cells treated with the potent AHR agonist FICZ (26) (Fig 1C). Moreover, the increased AHR expression observed in IDO1-expressing cells was not abolished upon treatment with AHR inhibitor CH223191 (27) (Fig 1D). Altogether, our results point to a direct regulation of AHR expression by the enzymatic activity of IDO1 and TDO. We next deciphered how the enzymatic activity of IDO1 and TDO regulates AHR expression.

### IDO1/TDO-induced AHR expression is induced by tryptophan deprivation

We first wanted to determine whether the increased AHR expression in IDO1-expressing cells was driven by kynurenine production and/or by tryptophan deprivation. When we treated wild-type HEK293-E cells for 72 hours with 80 μM fresh kynurenine, we observed no increase in AHR expression (Fig 2A). We then repeated this experiment adding kynurenine to the cells, but this time grown in tryptophan-free medium, to mimic the metabolic consequences of tryptophan catabolism mediated by IDO1 and TDO. We found no impact of kynurenine on AHR expression. Surprisingly, however, we found an increased AHR expression in cells cultured in the tryptophan-free medium (Fig 2A).

**Figure 2.**
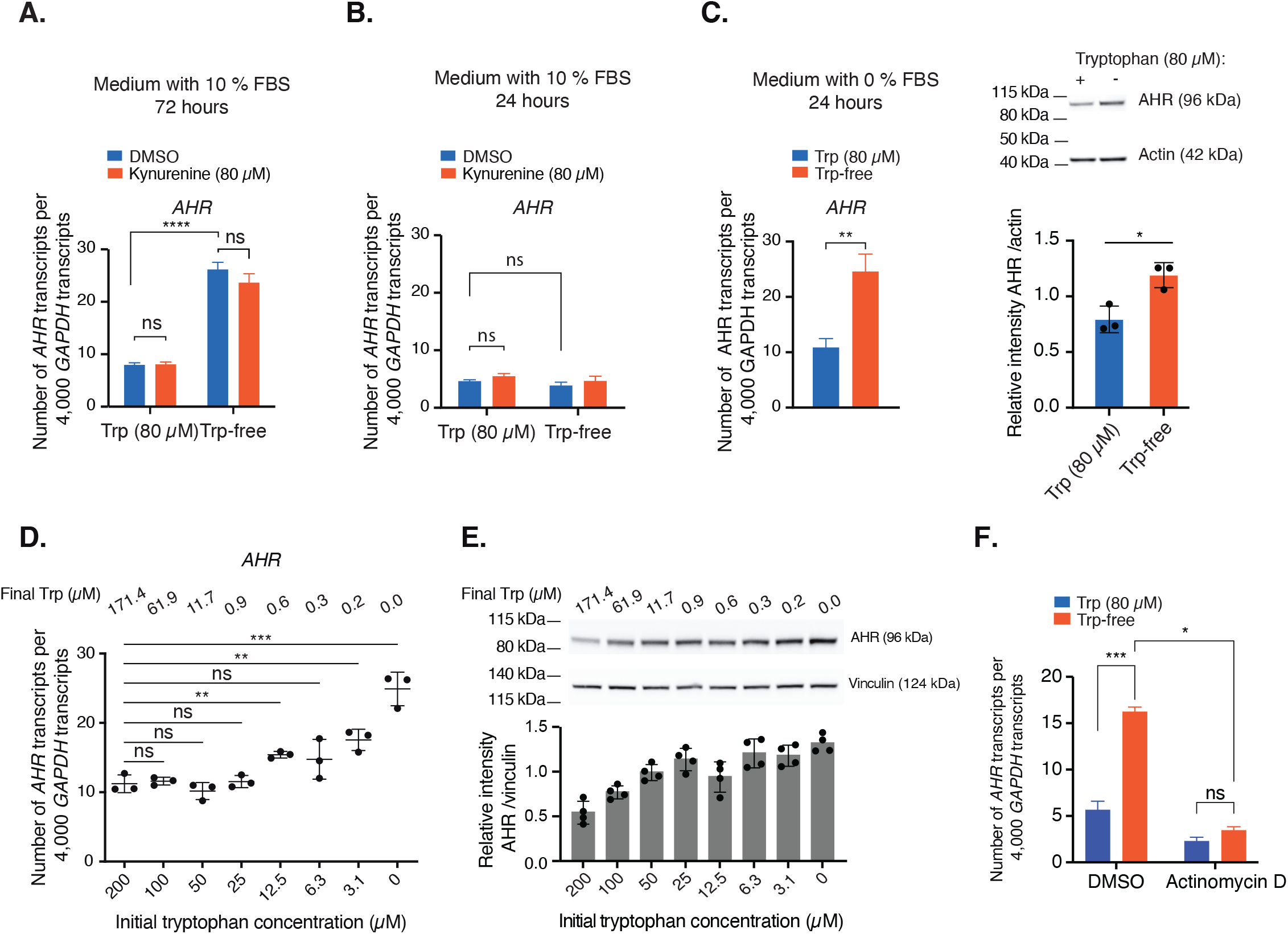
IDO1/TDO-induced AHR upregulation is induced by tryptophan deprivation. **A, B**. HEK293-E cells were treated with kynurenine (80 μM) in tryptophan-free medium supplemented or not with 80 μM tryptophan in the presence of 10 % FBS for 72 h **(A)** or 24 h **(B)**. *AHR* expression was measured by qRT-PCR analysis. Mean ± SD of technical triplicates from one representative experiment out of three independent experiments. HEK293-E cells were cultured for 24 h in tryptophan-free medium supplemented or not with 80 μM tryptophan in the absence of FBS. AHR expression was measured by qRT-PCR analysis (left panel) and western blot analysis (right panels). The quantification of AHR protein is shown in the lower right panel, each point representing a technical replicate. Mean ± SD of technical triplicates from one representative experiment out of three independent experiments (left panel). **D, E**. HEK293-E IDO1 cells were cultured in 2 % FBS with decreasing concentrations of tryptophan for 24 h. AHR expression was measured by qRT-PCR analysis **(D)** and western blot analysis **(E)**. Mean ± SD of technical triplicates from one representative experiment out of three independent experiments for panel D and Mean ± SD of technical duplicates from one representative experiment out of two independent experiments for panel E. In Figure 2E, the quantification of AHR is shown in the lower panel. Each point represents a technical replicate. **F**. HEK293-E cells treated with DMSO or Actinomycin D (5 µg/ml) were cultured for 24 h with or without tryptophan in medium without FBS. *AHR* expression was measured by qRT-PCR analysis. The mRNA levels of different genes were measured by quantitative qRT-PCR and normalized to *GAPDH*. Mean ± SD of technical triplicates from one representative experiment out of three independent experiments. **** significant with p<0.0001, *** significant with p<0.001, ** p<0.01, * p<0.05 and NS= not significant.

This result suggested that AHR expression was modulated by the tryptophan concentration. Increased AHR expression required very low tryptophan levels and was observed after 72h but not after 24 h of incubation in tryptophan-free medium (Fig 2A and 2B). This likely resulted from the time needed for the cells to completely consume the tryptophan coming with the fetal bovine serum (FBS) supplement. Indeed after 24 h there were about 2 µM residual tryptophan in the medium under these conditions (Suppl Fig 1B). Accordingly, when we cultured the cells in tryptophan-free medium without FBS, we observed an increased AHR expression already at 24 h (Fig 2C). To avoid the delayed effect in cells grown with FBS, we then used IDO1-overexpressing HEK293-E cells, which more actively degrade FBS-derived tryptophan. We incubated these cells for 24 h with different tryptophan concentrations, ranging from 0 to 200 µM, and evaluated AHR mRNA (Fig 2D) and proteins levels (Fig 2E). Tryptophan concentration in the supernatant was measured at the end of the incubation period (Supp Fig 1C). We observed an increase of AHR transcript and protein at low tryptophan levels. This effect was not restricted to HEK293-E cells, as an increase of *AHR* expression was also observed in colorectal carcinoma (LB159-CRC) and hepatocellular carcinoma (HepG2) cultured in tryptophan-depleted medium (Suppl Fig 2). Analysis of mRNA stability after transcriptional inhibition with Actinomycin D revealed that the increased *AHR* mRNA level in cells cultured under tryptophan deprivation was not due to increased mRNA stability but rather a direct regulation of *AHR* transcription (Fig 2F).

### AHR upregulation due to tryptophan depletion does not signal through GCN2

To further elucidate how tryptophan deprivation induces AHR expression, we assessed the involvement of GCN2, which senses unbound transfer RNA, indicating amino-acid shortage, and phosphorylates eukaryotic translation initiation factor 2α (eIF2α) to shut down cap-dependent translation (28), while favoring translation of activating transcription factor 4 (ATF4). The latter triggers transcription of genes involved in amino-acid metabolism

(29) and transportation (30), nutrient sensing and protein turnover (31, 32), in order to restore amino-acid homeostasis (29, 33). We confirmed that tryptophan depletion activates the GCN2/ATF4 pathway, as illustrated by increased phosphorylation of eIF2α and expression of ATF4 (Fig 3A). To test the involvement of this pathway, we first used salubrinal (34), an inhibitor of phosphatase “growth arrest and DNA damage-.inducible protein 34” (GADD34), which dephosphorylates eIF2α. Salubrinal increased ATF4 expression (Suppl Fig 3A), but did not increase AHR expression (Fig 3B). We also used Isrib (35), a compound that blocks the GCN2/eIF2α/ATF4 pathway downstream of eIF2α (Suppl Fig 3A) and observed no impact on the induction of AHR expression upon tryptophan depletion (Fig 3B). Finally, to confirm that the GCN2/eIF2α/ATF4 pathway was not involved in the regulation of AHR expression, we generated *GCN2*- and *ATF4*-knockout HEK293-E cells (Fig 3C) and assessed AHR expression in these cells under tryptophan-depleted conditions. As shown in Figures 3D and 3E, AHR expression was still increased upon tryptophan depletion, despite the absence of GCN2 and/or ATF4. Altogether, our results excluded the involvement of the GCN2/eIF2α/ATF4 pathway in the upregulation of AHR expression in response to tryptophan depletion.

**Figure 3.**
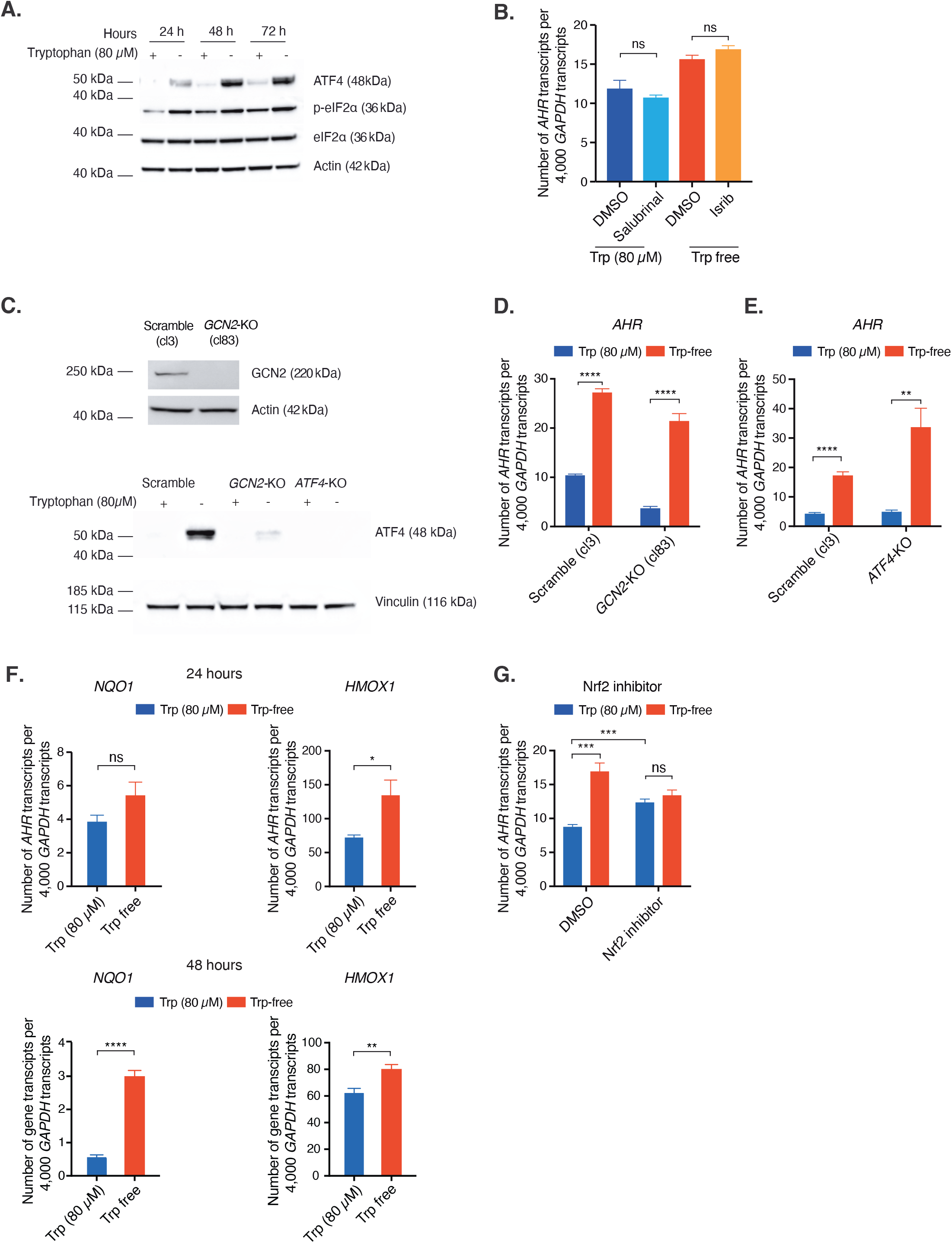
Tryptophan depletion-induced AHR overexpression is independent of GCN2. **A**. Western blot analysis of HEK293-E cells cultured with or without tryptophan for 24, 48 and 72 h for the expression of phospho-eIF2α, eIF2α, and ATF4. One representative experiment out of two independent experiments. **B**. qRT-PCR analysis of *AHR* expression in HEK293-E cells treated for 24 h with Salubrinal (20 μM), Isrib (5 μM) or vehicle (DMSO) with or without tryptophan in medium without FBS. Mean ± SD of technical triplicates from one representative experiment out of two independent experiments. **C**. Upper panel: *GCN2*-KO efficiency. Lower panel: ATF4 expression in *GCN2*-KO, *ATF4*-KO or wild-type cells cultured for 24 h without FBS in medium with or without tryptophan. One representative experiment out of two independent experiments. **D**. qRT-PCR analysis of *AHR* expression in scramble or *GCN2*-KO HEK293-E cells cultured for 24 h without FBS in medium with or without tryptophan. Mean ± SD of technical triplicates from one representative experiment out of three independent experiments. **E**. qRT-PCR analysis of *AHR* expression in wildtype or *ATF4*-KO HEK293-E cells cultured for 24 h without FBS in medium with or without tryptophan. Mean ± SD of technical triplicates from one representative experiment out of two independent experiments. **F**. qRT-PCR analysis of *NQO1* and *HMOX1* expression in HEK293-E cells cultured for 24 h or 48 h without FBS in medium with or without tryptophan. Mean ± SD of technical triplicates from one representative experiment out of three independent experiments. **G**. qRT-PCR analysis of *AHR* expression in HEK293-E cells cultured for 24 h without FBS in medium with or without tryptophan, and treated or not with NRF2 inhibitor, Brusatol (100 nM). The mRNA levels of different genes were measured by quantitative qRT-PCR and normalized to *GAPDH*. Mean ± SD of technical triplicates from one representative experiment out of two independent experiments. **** significant with p<0.0001, *** significant with p<0.001, ** p<0.01, * p<0.05 and NS= not significant.

Nuclear factor-erythroid factor 2-related factor 2 (NRF2) has been previously described to cross-talk with the AHR pathway. Earlier studies revealed that NRF2 could modulate the expression of AHR and vice versa (36, 37). An increase in NRF2 target genes was observed for HEK293-E cells under tryptophan deprivation, indicating an increase in NRF2 activation (Fig 3F). To know whether NRF2 regulates the increase in AHR transcription under tryptophan shortage, we treated cells with Brusatol, an NRF2 inhibitor that enhances the protein’s ubiquitination and degradation (38) (Suppl Fig 3B). As shown in Fig 3G, Brusatol treatment attenuated the increase of AHR transcription under tryptophan deprivation, suggesting the involvement of NRF2 in the increased AHR expression. Interestingly, we also observed an increased AHR expression when treating cells with Brusatol in the complete medium, indicating a role of other possible regulators in the crosstalk between NRF2 and AHR.

### AHR activation by kynurenine and derivatives is enhanced in the absence of tryptophan

Kynurenine has been widely described as a weak AHR agonist, although controversy remains due to its atypical, highly polar, and non-ligand-like structure (18). Given that tryptophan shortage increases AHR expression, we assessed whether it would potentiate AHR activation by kynurenine in HEK293-E cells, by analyzing the expression of the AHR target gene *CYP1A1*. We observed that, under normal tryptophan conditions, kynurenine did not activate AHR in these cells (Fig 4A). However, kynurenine clearly activated AHR in tryptophan-free conditions (Fig 4A), even at low concentrations (Fig 4B). The effect was AHR-dependent since adding an AHR inhibitor abolished the increase of expression of the AHR target gene under tryptophan deprivation (Fig 4A).

**Figure 4.**
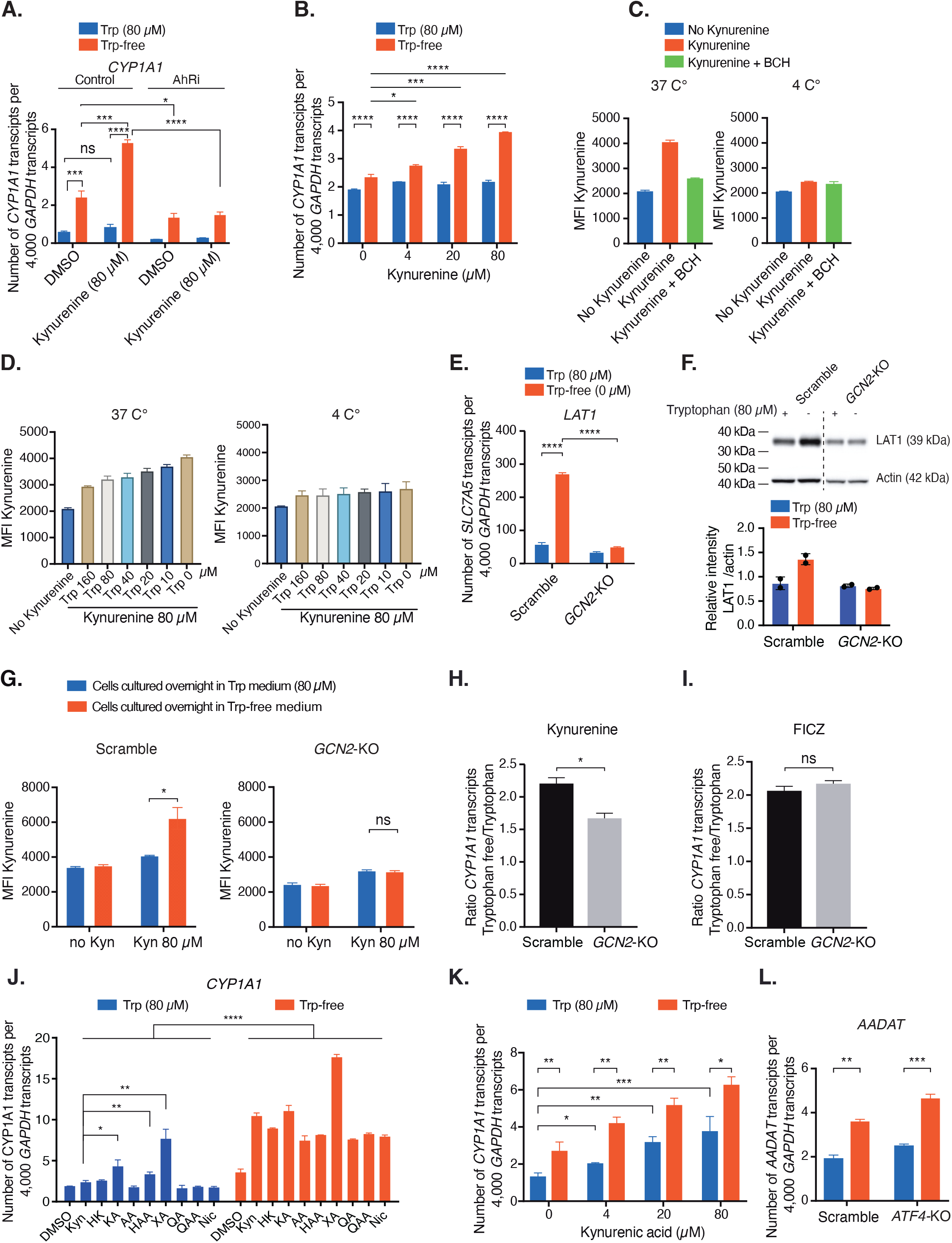
AHR activation by kynurenine and its derivatives is enhanced in the absence of tryptophan. **A**. HEK293-E cells were treated for 72 h with kynurenine (80 µM), AHR inhibitor (CH223191, 10 µM) or both in tryptophan-free medium supplemented or not with 80 µM tryptophan in the presence of 10 % FBS. *CYP1A1* expression was measured by qRT-PCR analysis. Mean ± SD of technical triplicates from one representative experiment out of three independent experiments. **B**. HEK293-E cells were treated for 24 h with different concentrations of kynurenine in tryptophan-free medium supplemented or not with 80 µM tryptophan in medium without FBS. *CYP1A1* expression was measured by qRT-PCR analysis. Mean ± SD of technical triplicates from one representative experiment out of two independent experiments **C**. Flow cytometric evaluation of kynurenine uptake in HEK293-E cells. Data acquired using 405LJnm excitation (violet laser) and band pass filter 525LJ±LJ50 on BD LSRII (Fortessa). Kynurenine fluorescence in HEK293-E cells treated with 80 µM kynurenine, in the presence or absence of the System L inhibitor, BCH (10LJmM) in PBS at 37 °C or 4 °C. **D**. Kynurenine fluorescence in HEK293-E cells treated with 80 µM kynurenine and different concentrations of tryptophan in PBS for 4 min. Mean ± SD of technical triplicates from one representative experiment out of three independent experiments. **E, F**. qRT-PCR analysis (**E**) and western blot analysis **(F)** of SLC7A5 expression in wildtype or *GCN2*-KO HEK293-E cells cultured in tryptophan medium or tryptophan-free medium. The quantification of LAT1 protein expression is shown in the lower panel, each point representing a technical replicate. Mean ± SD of technical triplicates from one representative experiment out of three independent experiments (panel E) and Mean ± SD of technical duplicates from one representative experiment out of two independent experiments (panel F). **G**. Scramble or *GCN2*-KO HEK293-E cells were cultured overnight in tryptophan-free medium supplemented or not with 80 µM tryptophan without FBS. Cells were then collected and incubated with 80 µM kynurenine during 4 min in PBS without tryptophan. Kynurenine uptake was measured by flow cytometry analysis of kynurenine fluorescence. Mean ± SD of technical triplicates from one representative experiment out of three independent experiments. **H, I**. qRT-PCR analysis of *CYP1A1* expression in wildtype or *GCN2*-KO HEK293-E cells treated for 24 h with kynurenine (80 µM) **(H)** or FICZ (1 µM) **(I)** in tryptophan-free medium supplemented or not with 80 µM tryptophan in the absence of FBS. Column graphs represent the fold increase of *CYP1A1* expression in cells cultured without tryptophan. The number of *CYP1A1* transcripts was first normalized to *GAPDH* before calculating the ratio between the tryptophan-free and the tryptophan condition. Mean ± SD of technical triplicates from one representative experiment out of three independent experiments. **J**. HEK293-E cells were treated for 72 h with 80 µM kynurenine metabolites in tryptophan-free medium supplemented or not with 80 µM tryptophan in the presence of 10 % FBS. *CYP1A1* expression was measured by qRT-PCR analysis. Mean ± SD of technical triplicates from one representative experiment out of three independent experiments. **K**. HEK293-E cells were treated for 24 h with different concentrations of kynurenic acid in medium with or without tryptophan and without FBS. *CYP1A1* expression was measured by qRT-PCR analysis. Mean ± SD of technical triplicates from one representative experiment out of two independent experiments **L**. qRT-PCR analysis of *AADAT* expression in wildtype or *ATF4*-KO HEK293-E cells cultured for 72 h in tryptophan-free medium supplemented or not with 80 µM tryptophan in the presence of 10 % FBS. The mRNA levels of different genes were measured by quantitative qRT-PCR and normalized to *GAPDH*. Mean ± SD of technical triplicates from one representative experiment out of two independent experiments. **** significant with p<0.0001, *** significant with p<0.001, ** p<0.01, * p<0.05 and NS= not significant.

Although in contrast with several reports in the literature (10-12), our observation that kynurenine does not activate AHR in normal conditions is in line with the report of Seok et al. indicating that kynurenine requires subsequent modifications to act as a potent AHR agonist (18). The authors showed that kynurenine stored in DMSO transforms into an oxidized dimeric derivative with a very potent AHR agonistic activity. We also observed a strong increase in the agonistic activity of our kynurenine solutions prepared in DMSO and identified the same derivative in our old solutions (Suppl Fig 4). These results suggest on one hand that some reports about AHR activation by kynurenine may have been affected by the presence of this strong artificial agonist. On the other hand, although there is no evidence that kynurenine can produce the same derivative “in cellulo”, it is possible that kynurenine acts as a “pro-ligand” requiring further metabolization to become a good AHR agonist. In our experiments, we used kynurenine freshly dissolved in medium, and we observed no AHR activation in normal conditions.

Note that there is a basal level of AHR activation in HEK293-E cells in the standard medium (Fig 4A), which might be due to the presence of aromatic AHR ligand-like compounds in the culture medium (39, 40). In line with this hypothesis, tryptophan depletion further increased this basal level of AHR activity, possibly due to increased AHR expression (Fig 4A). Similar observations were made for LB159-CRCA and HepG2 tumor cells (Suppl Fig 5).

Our results thus show that, under tryptophan deprivation, kynurenine is more potent to activate AHR, possibly due to increased AHR expression. Another factor that might potentiate the agonistic activity of kynurenine is its intracellular concentration. Cells have been reported to uptake tryptophan and kynurenine through the same System L transporter, SLC7A5 (LAT1) (41, 42). A decreased tryptophan concentration might thereby potentiate kynurenine uptake into the cells. To test this hypothesis, we adopted the flow cytometry-based assay developed previously by Sinclair et al. (42), which can directly monitor the intracellular level of kynurenine based on its fluorescence spectral properties. We further optimized the method and found a better signal detection when using a band pass filter 525/50 BP (Bv510) (Suppl Fig 6A, Fig 4C). As shown in Figure 4D, decreasing tryptophan concentration indeed potentiated kynurenine entry into the cells in a dose-dependent manner. In addition, we observed an increased expression of transporter SLC7A5 (LAT1) under tryptophan deprivation. The increase of LAT1 expression was regulated via the GCN2 pathway, as it was not observed in GCN2-KO cells (Fig 4E and F). These GCN2-KO cells also failed to increase kynurenine uptake in a tryptophan-free medium, indicating that this increase largely depended on LAT1 overexpression (Fig 4G, Suppl Fig 6B).

Tryptophan deprivation potentiates AHR activation by kynurenine in wild-type HEK293-E cells, as measured by CYP1A1 expression. This increase of AHR activation by kynurenine is partly due to the increased kynurenine entry mediated by the increased LAT1 expression upon tryptophan-deprivation, as a milder increase of *CYP1A1* is observed in GCN2-KO HEK293-E cells (Fig 4H). In comparison, the tryptophan-deprivation induced AHR overexpression plays a more prominent role since it potentiates AHR activation also by FICZ which does not depend on LAT1 for transportation (42) (Fig 4I). Altogether, our results indicate a multifaceted regulation of AHR activation by kynurenine under tryptophan deprivation, involving both GCN2-dependent and -independent pathways.

We also tested other kynurenine pathway metabolites and observed that tryptophan deprivation revealed a potent AHR agonistic activity for all of them (Fig 4J). Among them, kynurenic acid (KA) and xanthurenic acid (XA) showed already a more potent AHR agonistic activity in standard medium as compared to kynurenine. They became very potent in tryptophan-free medium (Fig 4J and K). To determine which kynurenine metabolite was produced in HEK293-E cells, we evaluated expression of the different enzymes of the pathway in these cells (Suppl Fig 6C). We found that kynurenine formamidase (*AFMID*) and kynurenine aminotransferase (*AADAT*) were expressed, but kynurenine 3-monooxygenase (*KMO*) and kynureninase (*KYNU*) were not. Therefore, the increased AHR activity in IDO1- or TDO-expressing HEK293-E cells or in kynurenine-treated HEK293-E cells cultured in tryptophan-free medium was possibly mediated by kynurenine and/or KA, but not by XA. Interestingly, we also observed a GCN2-ATF4 independent increased expression of kynurenine aminotransferase under tryptophan deprivation (Fig 4L). Altogether, the increased AHR activation by kynurenine under tryptophan deprivation might result from the combined effect of increased AHR expression, increased kynurenine import, and increased production of its metabolites.

### AHR upregulation favors *in vitro* differentiation of CD4^+^ FoxP3^+^ Treg cells

Analyzing TCGA patient tumor transcriptome data revealed a correlation between the expression of *AHR* and that of *IDO1/TDO2* in multiple tumor types, confirming the physiological relevance of our findings (Fig 5A). Among different immune cells in the TME, a strong correlation between AHR expression and the presence of Tregs was observed in various patient tumors (Fig 5B). It was previously observed that IDO1-expressing tumors had significantly more Tregs (43) and that the presence of AHR was required to optimally generate Treg cells *in vitro* (10). We therefore evaluated the role of tryptophan deprivation-induced AHR overexpression in Treg differentiation using murine wild-type and *Ahr*-KO mice. Two alleles encoding different *Ahr* variants exist among laboratory mouse strains (44) (Fig 5C). Compared to human AHR (hAHR), the mouse *Ahr*^b^ (m*Ahr*^b^) variant has a ∼10-fold higher affinity for TCDD (45). Most *in vivo* or *ex vivo* murine data published regarding the role of IDO1/TDO and AHR in immunosuppression result from experiments performed in C57BL/6 mice, which have the m*Ahr*^b^ variant. Using naïve CD4^+^ T cells derived from C57BL/6 WT or *Ahr*-KO mice, we confirmed the critical role of AHR for Treg differentiation induced by TGF-β *in vitro*. We observed that significantly less Tregs were generated by *Ahr*-KO as compared to WT CD4^+^ T cells (Fig 5D).

**Figure 5.**
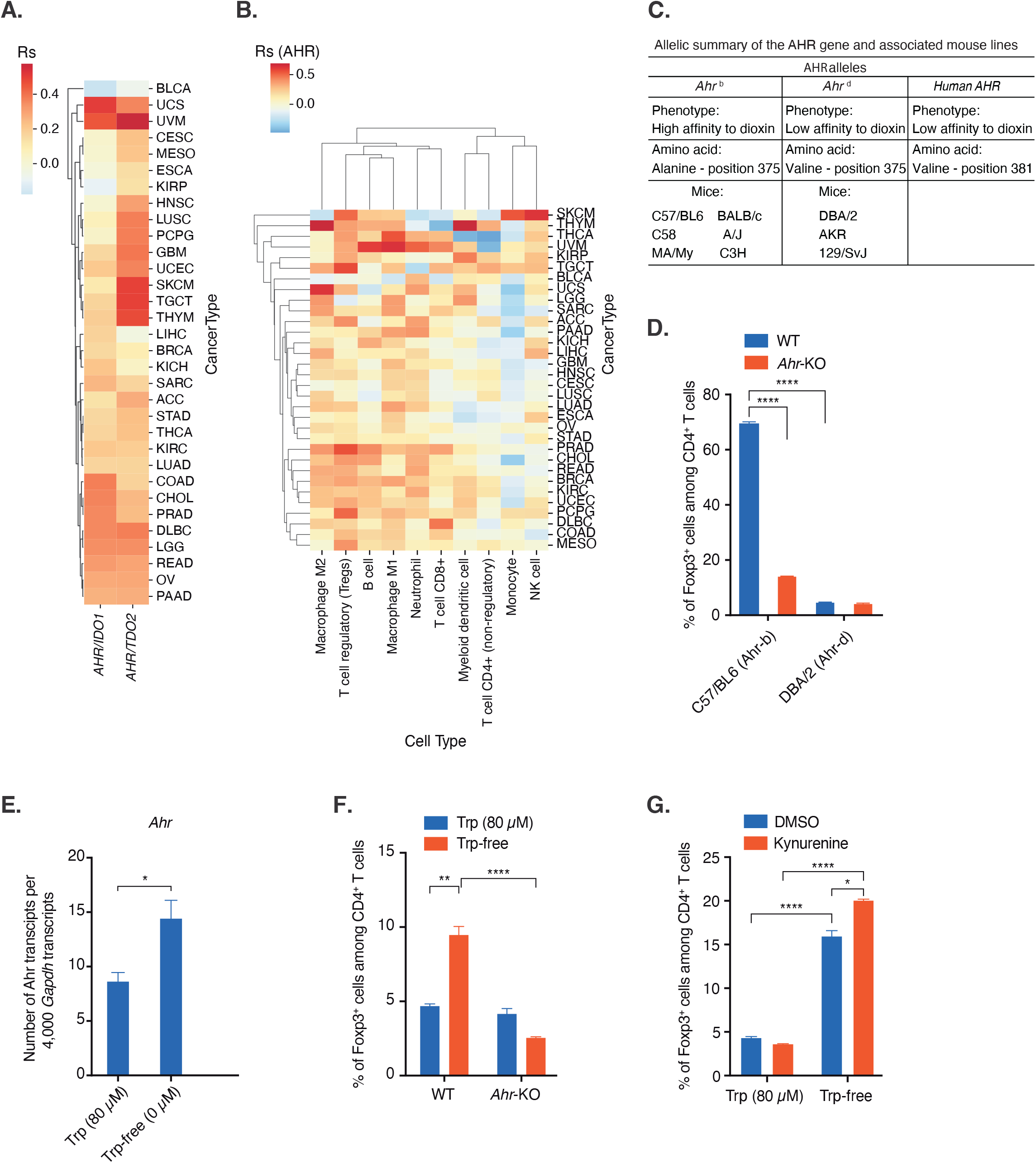
AHR upregulation favors *in vitro* differentiation of CD4^+^ Fox3^+^ regulatory T lymphocytes. **A**. Spearman correlation in bulk TCGA RNAseq data between expression of *AHR* and *IDO1* (left column) and *TDO2* (right column) per TCGA cancer type. **B**. Spearman correlation between expression of the *AHR* gene and various QUANTISEQ immune-deconvoluted populations per cancer type in the TCGA patient cohort. **C**. Allelic summary of the *AHR* gene in human and mouse lines. **D**. Naïve CD4^+^ CD62L^+^ T cells were isolated from WT (wild-type) and *Ahr*-KO mice from C57BL/6 or DBA/2 background and were cultured for 5 days with anti-CD3 anti-CD28 beads, TGF-β (5 ng/mL) and IL-2 (100 U/mL). The percentage of Treg differentiation was determined by flow cytometry analysis of cell surface CD4 and intracellular Foxp3 expression. **E**. qRT-PCR analysis of *Ahr* expression in Tregs differentiated from CD4^+^ T cells isolated from wild-type DBA/2 mice in medium containing or not 80 µM tryptophan. The mRNA levels of different genes were measured by quantitative qRT-PCR and normalized to *Gapdh*. Mean ± SD of triplicates from one out of three experiments. **F**. Treg differentiation (same as **D**) of naïve CD4^+^ CD62L^+^ T cells isolated from WT or *Ahr*-KO DBA/2 mice in medium containing or not 80 µM tryptophan. **G**. Treg differentiation (same as **D**) of naïve CD4^+^ CD62L^+^ T cells isolated from WT DBA/2 mice in medium containing or not 80 µM tryptophan in the presence or absence of 200 µM kynurenine. Mean ± SD of technical triplicates from one representative experiment out of three independent experiments. **** significant with p<0.0001, *** significant with p<0.001, ** p<0.01, * p<0.05 and NS= not significant.

DBA/2 mice carry the *Ahr*^d^ variant, which is structurally closer to h*AHR*, as it contains the same key valine residue in the active site (46). The m*Ahr*^d^ variant also has a lower affinity than m*Ahr*^b^ for TCDD, similar to that of hAHR (46). We observed that CD4^+^ T cells from DBA/2 mice produced very few Tregs after *in vitro* induction (Fig 5D). This might result from the low affinity of AHR^d^ for its ligands. When we repeated the experiment with DBA/2 CD4^+^ T cells in tryptophan-free medium, we observed an increased *AHR* expression (Fig 5E) and an enhanced Treg differentiation, which was abolished when we used CD4^+^ T cells from *Ahr*-KO/DBA/2 mice (Fig 5F). Treg differentiation was further enhanced upon addition of kynurenine, but only in tryptophan-free conditions (Fig 5G), in line with our previous observation that tryptophan deprivation potentiated AHR activation by kynurenine.

Altogether, our data confirm that AHR is required for Treg induction and indicate that tryptophan deprivation improves Treg induction by upregulating AHR expression. This new role of tryptophan deprivation appears to be key in CD4^+^ T cells that carry a low-affinity AHR variant, as is the case in DBA/2 mice, but also in humans.

## DISCUSSION

Our results show the impact of tryptophan deprivation on AHR regulation, demonstrating an increased expression of AHR at the transcript level, resulting in higher protein levels and a strong sensitization of the AHR pathway. A direct consequence is that weak AHR agonists, including kynurenine and its derivatives, become stronger agonists under tryptophan deprivation. Interestingly, in one of the first reports identifying kynurenine as an endogenous AHR agonist, kynurenine dissolved in aqueous solution was added to TDO-expressing cells cultured in tryptophan-free medium to study the effects of defined kynurenine concentrations on AHR activity by excluding endogenous production of kynurenine (12). In this study, it was already observed that the induction of the AHR target gene *CYP1A1* by kynurenine was stronger in cells cultured in tryptophan-free than in tryptophan-containing medium (12), but the underlying mechanisms remained elusive. The exact signaling pathway leading to AHR overexpression under tryptophan deprivation requires further elucidation, but our results suggest the involvement of the NRF2 pathway. This sensitization of the AHR pathway may contribute to a cellular response to overcome the stress induced by tryptophan depletion, in a process similar to the GCN2/ATF4 pathway reported by Fiore et al. (47). However, the two pathways are independent as AHR overexpression upon tryptophan deprivation does not depend on the GCN2/ATF4 pathway. Interestingly, in the same study (47), the authors observed that kynurenine pathway metabolites induced NRF2 expression.

Although its chemical structure does not make it an optimal AHR ligand, kynurenine has been proposed as an endogenous AHR agonist (12). It was also proposed that kynurenine could act as a proligand of AHR, requiring further metabolization to produce more potent agonist(s) (18). Seok et al. reported that kynurenine dissolved in DMSO spontaneously converts into aromatic condensation products that better fit the AHR ligand binding site and act as highly potent AHR agonists (18). We also observed that aged kynurenine solutions activated AHR much more strongly than fresh ones, and we identified the very same condensation derivative as a very potent AHR agonist in old kynurenine solutions. It is not clear at the moment whether this metabolite can be produced *in vivo* inside the cell. However, since kynurenine is often dissolved in DMSO, it is possible that the agonistic activity of kynurenine reported in the literature is partly due to this artificial contaminant.

When using fresh kynurenine solutions, we observed no AHR activation in HEK293-E cells. However, under tryptophan depletion, fresh kynurenine displayed AHR agonistic properties. This can result from several factors. Probably a dominant factor is the overexpression of AHR that we report here, which sensitizes the AHR pathway to kynurenine and its metabolites. A second factor could be the overexpression of LAT1, the transporter used by both tryptophan and kynurenine to enter cells. We show that contrary to AHR overexpression, LAT1 overexpression is dependent on the GCN2/ATF4 pathway. LAT1 overexpression results in increased import of kynurenine into the cell, potentiating its effect. Moreover, the mere absence of tryptophan should also favor kynurenine import, since tryptophan and kynurenine compete for the same transporter LAT1. Besides LAT1, other transporters of kynurenine have been described, although their impact on kynurenine entry in our experimental setting seems minor, as the LAT1 inhibitor BCH almost completely blocked kynurenine entry. It will, nevertheless, be interesting to evaluate the impact of tryptophan deprivation on other kynurenine transporters, such as SLC36A4, SLC7A8 (16), and SLC7A11 (47), using relevant cell lines. Lastly, we observed overexpression of kynurenine aminotransferase II under tryptophan deprivation in HEK293-E cells. This enzyme may increase production of kynurenine metabolites such as kynurenic acid, which may be more potent agonists than kynurenine itself.

Our results also point out the modulation of both amino acid and kynurenine uptake by cells in the tumor microenvironment, through changes in both amino-acid concentration and transporter expression. The latter is increased upon amino acid shortage in a GCN2-dependent manner (our results and Timosenko et al (30)). Furthermore, since kynurenine and hydrophobic amino acids both use LAT1 as an uptake transporter, kynurenine uptake is favored by the lack of tryptophan, at least in PBS that lack other amino acids. It remains to be determined whether the same is true in the TME, which contains other hydrophobic amino acids. Of note, local deprivation of other amino acids could also increase cellular uptake of kynurenine and tryptophan. Likewise, IDO1-expressing tumors produce high amounts of kynurenine, which could, in turn, affect the bioavailability of other amino acids transported by LAT1, and thereby regulate the function of immune cells in the tumor microenvironment. These interesting questions are worth further exploration.

The induction of AHR expression upon tryptophan deprivation has important consequences on immunoregulation. Indeed, we found that Treg differentiation induced by TGF-β was inefficient in DBA/2 mice in normal conditions but was strongly favored in tryptophan-depleted conditions. This was paralleled by an increased expression of *Ahr*. In *Ahr*-KO DBA/2 mice, differentiation of Tregs did not occur, even under tryptophan depletion. Altogether, this indicates that Treg differentiation requires a high expression of AHR to sensitize the pathway to weak agonists, and this can be obtained by tryptophan deprivation. In this protocol of *in vitro* Treg differentiation, there is no addition of exogenous kynurenine and no involvement of IDO1 (Suppl. Fig 7), the exact nature of the AHR ligand is unclear. It could be weak agonists present in the culture medium, as we observed (Fig 4A and S5) and reported in (39, 40). Addition of exogenous kynurenine further increased Treg differentiation in tryptophan-depleted conditions. Fallarino et al. reported already in 2006 that the induction of Treg *in vivo* by IDO1-expressing dendritic cells required both tryptophan depletion and kynurenine production (48). Yet no molecular explanation has been proposed so far for this observation. Our results provide a mechanistic explanation for this dual requirement for Treg induction: tryptophan depletion is needed to increase AHR and LAT1 expression, and thereby potentiate the weak AHR agonistic activity of kynurenine and derivatives. In the *in vivo* model of Fallarino et al., Treg induction was found to be dependent on GCN2, which suggests that the induction of LAT1 played a dominant role over the induction of AHR in these conditions.

Finally, we observed that the expression of AHR was a key limiting factor for Treg differentiation of CD4^+^ T cells from DBA/2 mice, which express the *Ahr*^d^ allele that has a low affinity for classical AHR ligands. CD4^+^ T cells from C57/BL6 mice, which express the high affinity allele *Ahr*^b^, do not need tryptophan depletion to differentiate into Tregs. The functional consequences of this allelic difference have been overlooked so far, as most studies analysing the role of AHR in immunoregulation made use of C57/BL6 mice. Importantly, the human AHR has the same valine residue as the murine AHR^d^ in the ligand binding site, and both have a low affinity for TCDD (46). Therefore, AHR results obtained in C57/BL6 mice should be extrapolated with caution to humans and might benefit from further validation in mice with an *Ahr*^d^ allele. Because our observation that Treg differentiation requires tryptophan depletion was made in DBA/2 mice, it is likely relevant to humans.

Ethics approval and consent to participate: Animal experimental procedures were approved by the Animal Ethical Committee of the UCLouvain (2015/UCL/MD/015; 2019/UCL/MD/019); the isolation of human blood cells was approved by the Comité d’Ethique Hospitalo-Facultaire Saint-Luc–UCLouvain.

## Consent for publication

All authors consent for publication

### Availability of data and material

All data relevant to the study are included in the article or uploaded as supplemental information. No datasets were generated in this study. All unique materials and reagents generated as part of this study are available from the lead contact with a completed Material Transfer Agreement.

### Funding

This work was supported by Ludwig Cancer Research, de Duve Institute (Belgium), and Université catholique de Louvain (Belgium). This work was also supported by grants from: Le Fonds de la Recherche Fondamentale Stratégique – WELBIO (Walloon Excellence in Life Sciences and Biotechnology), Belgium (grant number: WELBIO-CR-2019C-05); Fonds pour la Recherche Scientifique – FNRS, Belgium (grant number: EOS O000518F and PDR T.0091.18); Fondation contre le Cancer, Belgium (grant number: 2018-090); and the European Union’s Horizon 2020 Research and Innovation Programme (Grant Agreement No. 754688, MESI-STRAT (Systems Medicine of Metabolic-Signaling Networks) to CS, CO and BVdE. Fonds pour la Recherche Scientifique (FNRS), Belgium (Grant 23638468), Fédération Wallonie-Bruxelles (ARC 14/19-058). Q.S. was supported by a fellowship from FNRS-Télévie (grant 22846630). CO acknowledges support from the BMBF e:Med initiative GlioPATH (01ZX1402), the German Research Foundation (SFB1389 UNITE-Glioblastoma; project No. 404521405) and the European Research Council (ERC) under the European Union’s Horizon 2020 research and innovation programme (grant agreement number 101045257). PP was supported by the German Research Foundation (SFB1389 UNITE-Glioblastoma; project No. 404521405). JZ was supported by Fondation Contre le Cancer (Grant number 2019-094).

### Authors’contribution

MS, PP, SK, LP performed experiments, analysed data, and wrote materials methods and figure captions. VS designed and performed HPLC analysis. VS, JL, QS and RF designed and performed chemistry analysis of kynurenine derivative spontaneously produced in DMSO. SN designed and performed bioinformatics analyses. CS contributed to data interpretation and scientific discussions. EdP supervised the initial part of the study. CO analysed data and contributed to scientific discussions. JZ and BVDE supervised the study, designed and analysed experiments, and wrote the manuscript. All the authors read, revised, and approved the manuscript.

## Acknowledgments

We thank Pedro Gomez and the Platform Laboratory Animal Facility for mouse colony production, Isabelle Grisse for editorial assistance. Mihn-Phuong Le and Loubna Boudhan for the technical assistance. The results shown in this study are in whole or part based upon data generated by the TCGA Research Network: https://www.cancer.gov/tcga.

## List of abbreviations

3-HK: 3-Hydroxykynurenine
2-ME: 2-Mercaptoethanol
AA: Anthranilic acid
AFMID: Aryl formamidase
AHR: Aryl hydrocarbon receptor
ATF4: Activating transcription factor 4
BCA: Bicinchoninic acid assay 2-Aminobicyclo[2.2.1]heptane-2-carboxylic
BCH: acid
BSA: Bovine serum albumin
C57/BL6: C57/ Black 6 mice
CD62-L: L-selectin
CD8: Cluster of differentiation 8
CYP1-A1,: -B1, A2 Cytochrome P450
DBA/2: Diluted Brown Non-Agouti/2
DMSO: Dimethyl sulfoxide
DNA: Deoxyribonucleic acid
eIf2α: Eukaryotic translation initiation factor 2 alpha
EREG: Epiregulin
FBS: Fetal bovine serum
FICZ: 6-formylindolo (3,2-b) carbazole
FoxP3: Forkhead box protein P3
GCN2: General control nonderepressible 2
HAA: 3-OH anthranilic acid
HBSS: Hanks’ Balanced Salt Solution
HepG2: Heptacarcinoma cell line (human)
HEK293-E: Human embryonic kidney 293 cells HK 3-OH kynurenine
HPLC: High-performance liquid chromatography
HRP: Horse radish peroxydase
IDO1: Indoleamine 2,3-dioxygenase 1
IL-10: Interleukin 10
IL4I1: Interleukin 4 induced 1
KA: Kynurenic acid
KAT2: (aadat) Kynurenine aminotransferase KMO Kynurenine monooxygenase
KO: Knockout
Kyn: Kynurenine
KYNU: Kynureninase
LB159-CRC: Colorectal carcinoma cell line (human) LAT1 Large amino acid transporter 1
LT-K: Low tryptophan and kynurenine
MOPS: 3-(N-morpholino)propanesulfonic acid
NIC: Nicotinamide
NP40: Nonyl phenoxypolyethoxylethanol
PBS: Phosphate-buffered saline
PD-1: Programmed cell death protein 1
QA: 2,3-pyridinedicarboxylic acid
QAA: Quinaldic acid
RNA: Ribonucleic acid
SDS: Sodium dodecyl sulfate
STC2: Stanniocalcin 2
TAM: Tumor associated macrophage
TBS: Tris buffered saline
TCDD: 2,3,7,8 tetrachlorodibenzo-p-dioxin
TDO: Tryptophan 2,3-dioxygenase
TGF-β: Transforming growth factor-b
TIPARP: TCDD Inducible Poly(ADP-Ribose) Polymerase
TME: Tumor microenvironment
Treg: Regulatory T cell
tRNAs: Transfer RNAs
UV: Ultra Violet
XA: Xanthurenic acid

## FIGURE LEGENDS

**Supplementary Figure 1: IDO1 and TDO activity induces AHR activation**

**A**. qRT-PCR analysis of *CYP1B1, TIPARP, STC2, EREG* and *IL4I1* expression in HEK293-E or HEK293-E hIDO1 cells treated or not with IDO1 inhibitor (Epacadostat, 2.5 µM) for 120 h. The mRNA levels of different genes was measured by quantitative qRT-PCR and normalized to *GAPDH*. Mean ± SD of technical triplicates from one representative experiment out of three independent experiments. **** significant with p<0.0001, *** significant with p<0.001, ** p<0.01, * p<0.05 and NS= not significant. **B. T**ryptophan concentrations measured by HPLC in cell supernatant from Fig **2A** and **2B. C**, HPLC quantification of kynurenine or tryptophan concentrations in the supernatant of HEK293-E IDO1 cells from the experiment shown in Fig **2D** and **E**.

**Supplementary Figure 2: Tryptophan deprivation induces AHR expression in human tumor cells of different histological types**

Colorectal carcinoma (LB159-CRC) and hepatocellular carcinoma (HepG2) cells were cultured for 72 h in tryptophan-free medium supplemented or not with 80 µM tryptophan, in the presence of 10 % FBS. AHR expression was measured by quantitative qRT-PCR analysis and normalized to *GAPDH*. Mean ± SD of technical triplicates from one representative experiment out of three independent experiments. **** significant with p<0.0001, *** significant with p<0.001, ** p<0.01, * p<0.05 and NS= not significant.

**Supplementary Figure 3: Western blot analysis of the integrated stress response (ISR) in cells treated with Salubrinal or Isrib**

**A**. Western blot analysis of the expression of ATF4 in HEK293-E cells treated for 24 h with Salubrinal (20 µM), Isrib (5 µM) or vehicle (DMSO) in tryptophan-free medium supplemented or not with 80 µM tryptophan in the absence of FBS. One representative out of three independent experiments. **B**. Western blot analysis of the expression of NRF2 in HEK293-E cells treated for 24 h with brusatol (100 nM) or vehicle (DMSO). One representative out of two independent experiments.

**Supplementary Figure 4: A kynurenine derivative spontaneously produced in DMSO potently activates AHR**

Kynurenine is commonly dissolved in DMSO. However, in DMSO kynurenine is transformed into an oxidized dimeric derivative that potently activates AHR. Here under we refer to the transformed kynurenine as «Old» kynurenine (stored at RT for at least 5 days) and freshly dissolved kynurenine as «New».

**A.** HepG2 cells were treated with 4 different concentrations of each compound: 100 μM, 25 μM, 5 μM, 1 μM of kynurenine «Old» (•) and kynurenine «New»(▪). Known AHR ligands 2,3,7,8 TCCD (*) and FICZ (□) were used as controls at 30 nM and at 1 μM, 250 nM, 50 nM and 10 nM respectively. *CYP1A1* expression was measured by qRT-qPCR after 24 h (left). Data presented are the means ± SD and are representative of 3 independent experiments. AHR activation was measured in a luciferase assay after 16 h (right). A construct containing 4 repeats of the target sequence of AHR followed by the sequence of the reporter gene firefly luciferase was stably transfected in HepG2 cells (HepG2 PGudLuc6.1). Data presented are the means ± SD and are representative of 3 independent experiments. *** significant with p<0.001, ** p<0.01, * p<0.05 and NS= not significant.

**B.** The two kynurenine solutions were analysed by HPLC on a C18 column. New peaks were detected by UV in the «Old» solution in addition to the two major peaks corresponding to DMSO and kynurenine, while the kynurenine peak was reduced by a factor 1.7 in the «Old» solution. This indicated the presence of other molecules than kynurenine in the «Old» solution.

**C.** To identify the molecule responsible for activating AHR in the «Old» solution, a HPLC separation system was used to isolate forty fractions from both the «Old» (white) and the «New» (black) solutions. These fractions were tested in an AHR luciferase assay as described in A. Control (CTRL) corresponds to the unfractionated kynurenine solution. These data are representative of 3 independent experiments. The intact kynurenine present in fraction 3-4 had no activity. Fraction 28 was the most active in 3 independent experiments. The blue and red lines represent the UV spectrum of «New» and «Old» kynurenine, respectively.

**D.** Putative structure of the compound (K270) that activates AHR in fraction 28 based on kynurenine structure and chemical analysis.

**E.** Electrospray ionization (ESI-MS) analysis of «Old» kynurenine fraction 28 provided accurate mass measurement of the ion [M+H]^+^ at m/z 270.1027, and the molecule was therefore named K270. Based on the structure prediction, the compound was synthesised following a four-steps procedure involving the initial formation of the ethyl 4-methyl-2-oxo-1,2-dihydroquinoline-3-carboxylate by condensation and cyclisation of the commercially available 2-aminoacetophenone with the ethyl 3-chloro-3-oxopropanoate. The cyclized compound was next chlorinated with oxalyl chloride to afford the ethyl 2-chloro-4-methylquinoline-3-carboxylate derivative. In the next step, the tetracyclic 12-methyldibenzo[*b,g*][1,8]naphthyridin-11(6*H*)-one was obtained by reaction of the latter with aniline and polyphosphoric acid (PPA). The final 9*H*-benzo[*h*]quinolino[4,3,2-*de*][1,6]naphthyridine compound was obtained by reaction of this tetracyclic precursor with Bredereck’s reagent. It should be noted that the final compound (QS006) was only obtained in poor yield because of the difficulty of the purification step that could only be performed by isolation of the pure product after low-efficient HPLC separations. Figure shown is the spectroscopy analysis of K270 and QS006 (synthesized), with perfectly superimposed results confirming that QS006 (blue line) is identical or highly similar to K270 (red line).

**F.** The two molecules K270 and QS006 were tested for their ability to activate AHR under normal tryptophan (80 µM) containing conditions in a Luciferase reporter assay described in **A**. TCDD was used as a control for AHR activation. NT: not treated.

**G**. HepG2 cells were incubated with K270 or QS006 at two concentrations (78 nM and 39 nM) for 24 h and the induction of *CYP1A1* was measured by qRT-PCR. TCDD was used as a control for AHR activation.

Data presented are the Mean ± SD of technical triplicates from one representative experiment out of three independent experiments. *** significant with p<0.001, ** p<0.01, * p<0.05 and NS= not significant.

**Supplementary Figure 5: Tryptophan deprivation induces AHR activation in human tumor cells of different types**

Colorectal carcinoma LB159-CRC cells or hepatocarcinoma HepG2 cells were cultured for 72 h in tryptophan-free medium supplemented or not with 80 μM tryptophan in the presence of 10 % FBS. *CYP1A1* and *TIPARP* expression were measured by qRT-PCR analysis. The mRNA levels of different genes were measured by quantitative qRT-PCR and normalized to *GAPDH*. Mean ± SD of technical triplicates from one representative experiment out of three independent experiments. **** significant with p<0.0001, *** significant with p<0.001, ** p<0.01, * p<0.05 and NS= not significant.

**Supplementary Figure 6: Tryptophan deprivation induces LAT1 expression via the GCN2 pathway**

**A**. Flow cytometric evaluation of kynurenine uptake in HEK293-E cells. HEK293-E cells were treated with different concentrations of kynurenine in PBS for 4 minutes at 37 C°. The reaction was stopped after 4 minutes by adding 125 μl 4 % PFA for 30 minutes at room temperature in the dark. Data was acquired using 405 nm excitation (violet laser) and bandpass filter 525/50 BP (Bv510) on BD LSRII (Fortessa). **B**. Western blot analysis of LAT1 expression for Fig **4G**.

**C**. qRT-PCR analysis of the expression of enzymes *AFMID (kynurenine formamidase), AADAT (kynurenine aminotransferase), KMO (kynurenine 3-monooxygenase)*, and *KYNU (kynureninase)*, which are involved in kynurenine metabolism, in HEK293-E cells. Mean ± SD of technical triplicates from one representative experiment out of three independent experiments. **** significant with p<0.0001, *** significant with p<0.001, ** p<0.01, * p<0.05 and NS= not significant.

**Supplementary Figure 7: Impact of IDO on CD4**^**+**^ **FoxP3**^**+**^ **Treg-cell differentiation**

Treg differentiation (same as Fig 5D) of naïve CD4^+^ CD62L^+^ T cells isolated from WT mice treated or not with IDO1 inhibitor (Epacadostat, 2.5µM) during Treg differentiation. Treg differentiation (same as Fig 5D) of naïve CD4^+^ CD62L^+^ T cells isolated from WT or *Ido1*-KO mice. Mean ± SD of technical triplicates from one representative experiment out of three independent experiments. **** significant with p<0.0001, *** significant with p<0.001, ** p<0.01, * p<0.05 and NS= not significant.

## SUPPLEMENTARY METHODS

### Transfections and Luciferase Assays

The pGudLuc6.1 plasmid containing a mouse MMV viral promoter and 4 DRE binding sites upstream of a firefly luciferase reporter was a kind gift of Jing Zhao and Michael S. Denison from the department of Environmental Toxicology, University of California (49, 50). The plasmid was stably co-transfected in the HepG2 cell line with pGKNeo (conferring resistance to neomycin) using ExGen 500 transfection reagent from Fermentas. 5mg of the pGKNeo plasmid and 20mg of the pGudLuc6.1 plasmid were mixed in 1ml of NaCl 150mM. 82.5ml of ExGen 500 was added and then vortexed during 10 seconds. After 10 minutes of incubation at room temperature, the mixture was added gently in Petri dishes (100mm) on cells at around 80 % confluence. After 40 h, cells were split into 24-well plates and 800 mg/ml of G418 (Roche) were added for selection of positive populations (Protocol adapted from Long *et al*. 1998 (49)). For the luciferase assay, 50,000 cells per well were plated in 96-well plates. After 24 h, 100 mM, 25 mM, 5 mM and 1mM of tryptophan or tryptophan metabolites, 1 µM, 250 nM, 50 nM, 10 nM of FICZ or 30 nM, 10 nM, 3 nM and 1 nM of 2,3,7,8 TCDD were added to the cells. 16 h later, the luciferase activity of the cells was measured using the Luciferase reagent from Promega (Dual-Luciferase ® Reporter Assay System). Results were normalised on the protein concentration measured with the BCA protein assay kit (Pierce™).

### HPLC, UV analysis, mass spectrometry and NMR

UV spectra were recorded on an Ultrospec™ 2100 pro UV/Visible Spectrophotometer (GE Healthcare Life Sciences). HPLC analyses were performed with a Waters 616 pump equipped with a Waters 996 PDA detector and a Waters 717 autosampler. Kynurenine “old” and “new” (Fig 3B and C) were analysed by reverse-phase chromatography on a 3.9 × 150-mm Delta Pack C18 column (Waters) at a flow rate of 0.9 mL/min with a 35-min linear gradient of 3–75 % acetonitrile (ACN) in water, both with 0.1 % trifluoroacetic acid (TFA) (wt/vol). Kynurenine “old” and “new” (for the luciferase assay-Fig 3D and 4E) were further fractionated by HPLC with an elution gradient of 5–30 % acetonitrile/0.1 % TFA over 35 min followed by 30–80 % acetonitrile / 0.1 % TFA over 5 min. Fractions of 0.9 mL were collected, dried in a centrifugal vacuum concentrator, and dissolved in a solution of PBS-DMSO 15 %. The mass spectrometry analysis was performed using a LCQ Deca XP ion-trap spectrometer equipped with an electrospray ionization source (ThermoFinnigan). The sample dissolved in methanol was introduced directly into the source at a flow rate of 4 μl/minute. The LCQ was operated in positive mode under manual control in the Tune Plus view with default parameters and active automatic gain control. Accurate mass measurements were obtained using an LTQ Orbitrap mass spectrometer (ThermoFisher Scientific).

