## Supplementary Figures for "Tryptophan depletion sensitizes the AHR pathway by increasing AHR expression and GCN2/LAT1-mediated kynurenine uptake, and potentiates induction of regulatory T lymphocytes"

Supplementary Figure 1

**A.**

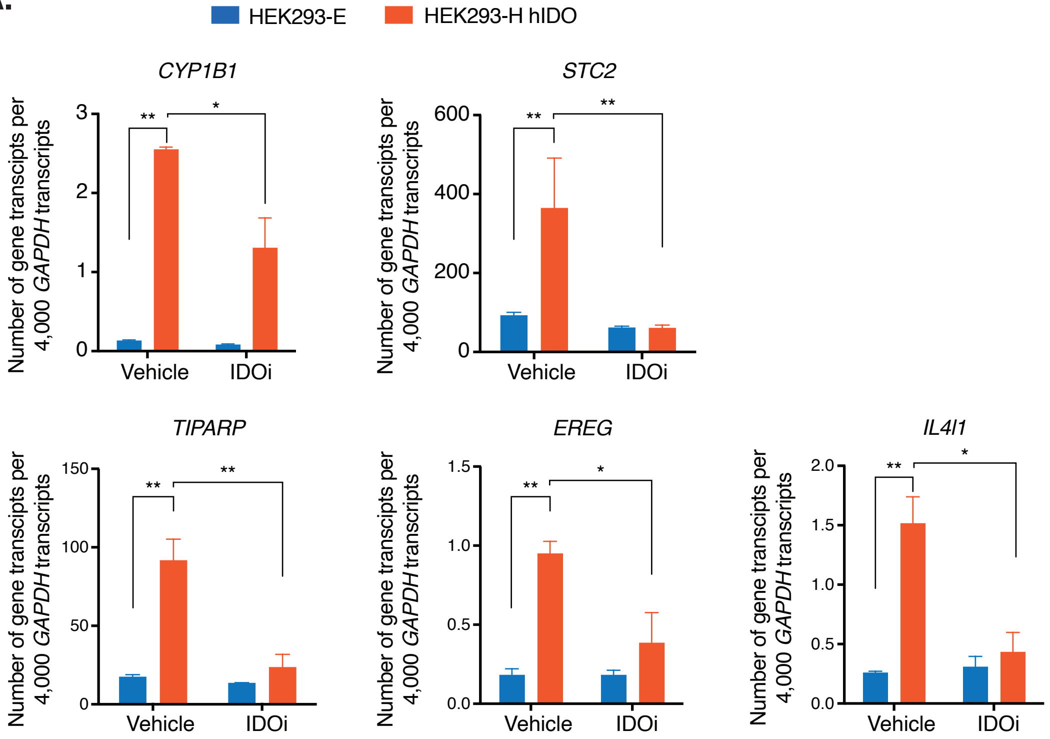


**B.**

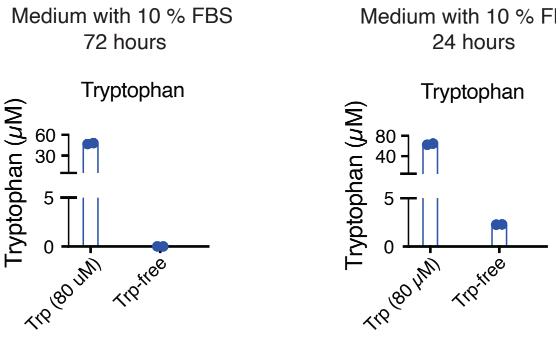


**C.**

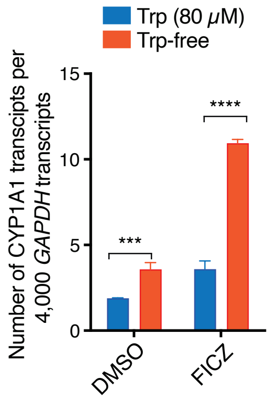


**D.**

| Initial Trp. Conc (µM) | Final Trp. Conc (µM) | Final Kyn. Conc (µM) |
|---|---|---|
| 200.0 | 171.4 | 31 |
| 100.0 | 61.9 | 31.1 |
| 50.0 | 11.7 | 31.4 |
| 25.0 | 0.9 | 16.4 |
| 12.5 | 0.6 | 8.4 |
| 6.3 | 0.3 | 5.5 |
| 3.1 | 0.2 | 2.9 |
| 0.0 | 0.0 | 1.3 |

**Supplementary Figure 1: IDO1 and TDO activity induces AHR activation**
**A.** qRT-PCR analysis of *CYP1B1, TIPARP, STC2, EREG* and *IL4I1* expression in HEK293-E or HEK293-E hIDO1 cells treated or not with IDO1 inhibitor (Epacadostat, 2.5 µM) for 120 h. The mRNA levels of different genes was measured by quantitative qRT-PCR and normalized to *GAPDH*. Mean + SD of technical triplicates from one representative experiment out of three independent experiments. **** significant with p<0.0001, *** significant with p<0.001, ** p<0.01, * p<0.05 and NS= not significant. **B.** Tryptophan concentrations measured by HPLC in cell supernatant from Fig **2A** and **2B**. **C**, HPLC quantification of kynurenine or tryptophan concentrations in the supernatant of HEK293-E IDO1 cells from the experiment shown in Fig **2D** and **E**.

Supplementary Figure 2

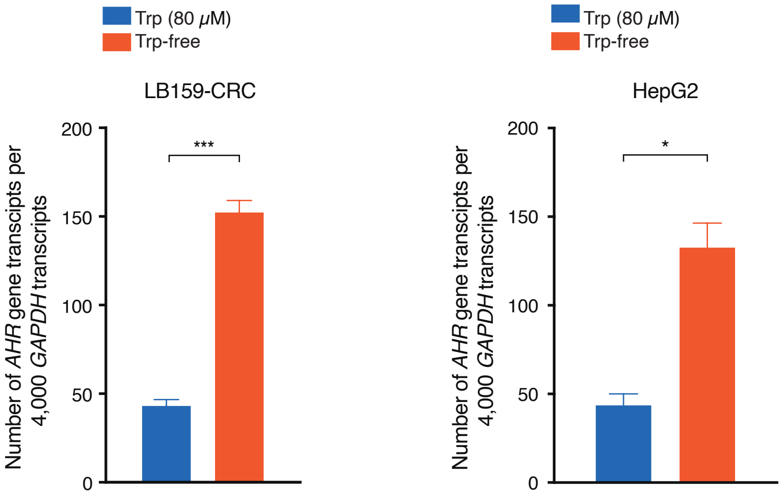


**Supplementary Figure 2: Tryptophan deprivation induces AHR expression in human tumor cells of different histological types**

Colorectal carcinoma (LB159-CRC) and hepatocellular carcinoma (HepG2) cells were cultured for 72 h in tryptophan-free medium supplemented or not with 80 µM tryptophan, in the presence of 10 % FBS. AHR expression was measured by quantitative qRT-PCR analysis and normalized to *GAPDH*. Mean + SD of technical triplicates from one representative experiment out of three independent experiments. **** significant with $p<0.0001$, *** significant with $p<0.001$, ** $p<0.01$, * $p<0.05$ and NS= not significant.

Supplementary Figure 3

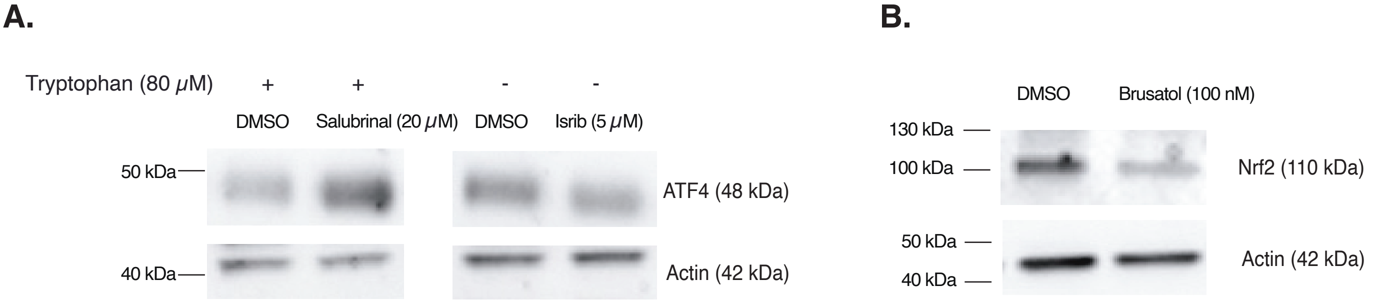


**Supplementary Figure 3: Western blot analysis of the integrated stress response (ISR) in cells treated with Salubrinal or Isrib**

**A.** Western blot analysis of the expression of ATF4 in HEK293-E cells treated for 24 h with Salubrinal (20 μM), Isrib (5 μM) or vehicle (DMSO) in tryptophan-free medium supplemented or not with 80 μM tryptophan in the absence of FBS. One representative out of three independent experiments. **B.** Western blot analysis of the expression of NRF2 in HEK-293-E cells treated for 24 h with brusatol (100 nM) or vehicle (DMSO). One representative out of two independent experiments.

Supplementary Figure 4

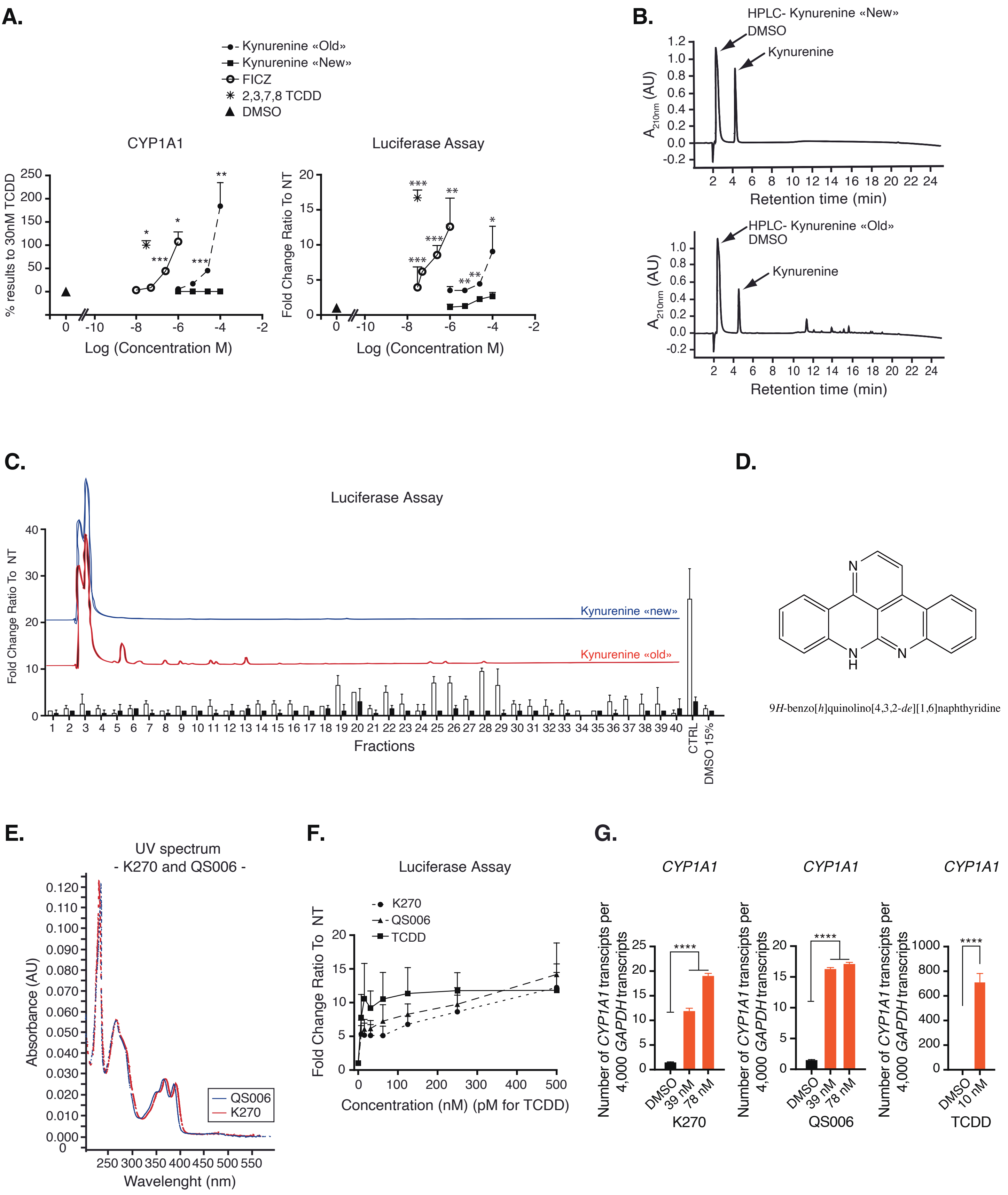

**Supplementary Figure 4: A kynurenine derivative spontaneously produced in DMSO potently activates AHR**
Kynurenine is commonly dissolved in DMSO. However, in DMSO kynurenine is transformed into an oxidized dimeric derivative that potently activates AHR. Here under we refer to the transformed kynurenine as «Old» kynurenine (stored at RT for at least 5 days) and freshly dissolved kynurenine as «New».

**A.** HepG2 cells were treated with 4 different concentrations of each compound: 100 µM, 25 µM, 5 µM, 1 µM of kynurenine «Old» (●) and kynurenine «New»(■). Known AHR ligands 2,3,7,8 TCCD (✱ ) and FICZ (□) were used as controls at 30 nM and at 1 µM, 250 nM, 50 nM and 10 nM respectively. *CYP1A1* expression was measured by qRT-qPCR after 24 h (left). Data presented are the means ± SD and are representative of 3 independent experiments. AHR activation was measured in a luciferase assay after 16 h (right). A construct containing 4 repeats of the target sequence of AHR followed by the sequence of the reporter gene firefly luciferase was stably transfected in HepG2 cells (HepG2 PGudLuc6.1). Data presented are the means ± SD and are representative of 3 independent experiments. *** significant with $p<0.001$, ** $p<0.01$, * $p<0.05$ and NS= not significant.

**B.** The two kynurenine solutions were analysed by HPLC on a C18 column. New peaks were detected by UV in the «Old» solution in addition to the two major peaks corresponding to DMSO and kynurenine, while the kynurenine peak was reduced by a factor 1.7 in the «Old» solution. This indicated the presence of other molecules than kynurenine in the «Old» solution.

**C.** To identify the molecule responsible for activating AHR in the «Old» solution, a HPLC separation system was used to isolate forty fractions from both the «Old» (white) and the «New» (black) solutions. These fractions were tested in an AHR luciferase assay as described in A. Control (CTRL) corresponds to the unfractionated kynurenine solution. These data are representative of 3 independent experiments. The intact kynurenine present in fraction 3-4 had no activity. Fraction 28 was the most active in 3 independent experiments. The blue and red lines represent the UV spectrum of «New» and «Old» kynurenine, respectively.

**D.** Putative structure of the compound (K270) that activates AHR in fraction 28 based on kynurenine structure and chemical analysis.

**E.** Electrospray ionization (ESI-MS) analysis of «Old» kynurenine fraction 28 provided accurate mass measurement of the ion $[M+H]^+$ at m/z 270.1027, and the molecule was therefore named K270. Based on the structure prediction, the compound was synthesised following a four-steps procedure involving the initial formation of the ethyl 4-methyl-2-oxo-1,2-dihydroquinoline-3-carboxylate by condensation and cyclisation of the commercially available 2-aminoacetophenone with the ethyl 3-chloro-3-oxopropanoate. The cyclized compound was next chlorinated with oxalyl chloride to afford the ethyl 2-chloro-4-methylquinoline-3-carboxylate derivative. In the next step, the tetracyclic 12-methyldibenzo[*b,g*][1,8]naphthyridin-11(6*H*)-one was obtained by reaction of the latter with aniline and polyphosphoric acid (PPA). The final 9*H*-benzo[*h*]quinolino[4,3,2-*de*][1,6]naphthyridine compound was obtained by reaction of this tetracyclic precursor with Bredereck's reagent. It should be noted that the final compound (QS006) was only obtained in poor yield because of the difficulty of the purification step that could only be performed by isolation of the pure product after low-efficient HPLC separations. Figure shown is the spectroscopy analysis of K270 and QS006 (synthesized), with perfectly superimposed results confirming that QS006 (blue line) is identical or highly similar to K270 (red line).

**F.** The two molecules K270 and QS006 were tested for their ability to activate AHR under normal tryptophan (80 *µ*M) containing conditions in a Luciferase reporter assay described in A. TCDD was used as a control for AHR activation. NT: not treated.

**G.** HepG2 cells were incubated with K270 or QS006 at two concentrations (78 nM and 39 nM) for 24 h and the induction of *CYP1A1* was measured by qRT-PCR. TCDD was used as a control for AHR activation.

Data presented are the Mean + SD of technical triplicates from one representative experiment out of three independent experiments.*** significant with $p<0.001$, ** $p<0.01$, * $p<0.05$ and NS= not significant.

Supplementary Figure 5

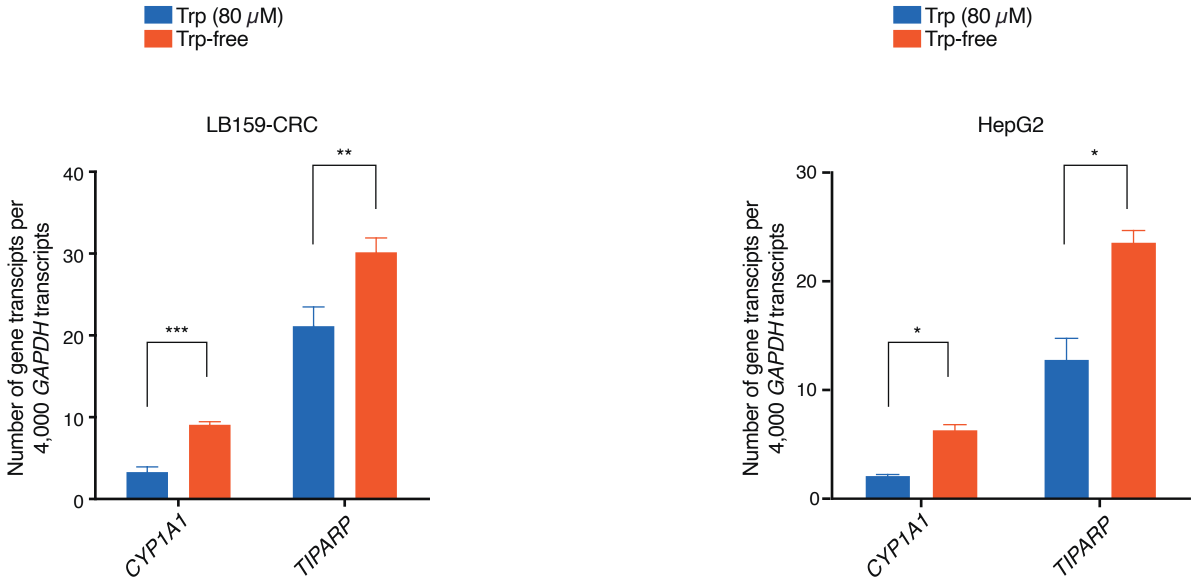


**Supplementary Figure 5: Tryptophan deprivation induces AHR activation in human tumor cells of different types**

Colorectal carcinoma LB159-CRC cells or hepatocarcinoma HepG2 cells were cultured for 72 h in tryptophan-free medium supplemented or not with 80 µM tryptophan in the presence of 10 % FBS. *CYP1A1* and *TIPARP* expression were measured by qRT-PCR analysis. The mRNA levels of different genes were measured by quantitative qRT-PCR and normalized to *GAPDH*. Mean + SD of technical triplicates from one representative experiment out of three independent experiments. **** significant with p<0.0001, *** significant with p<0.001, ** p<0.01, * p<0.05 and NS= not significant.

Supplementary Figure 6

A.

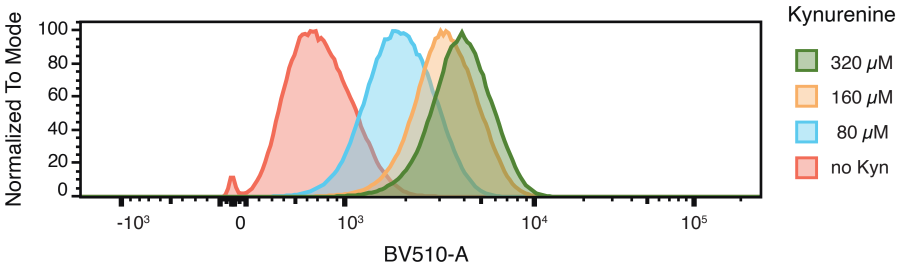


B.

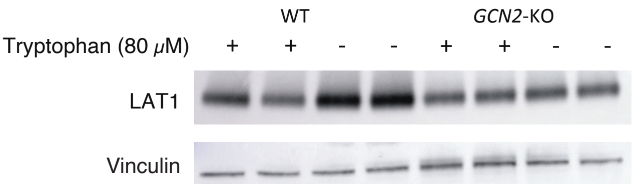


C.

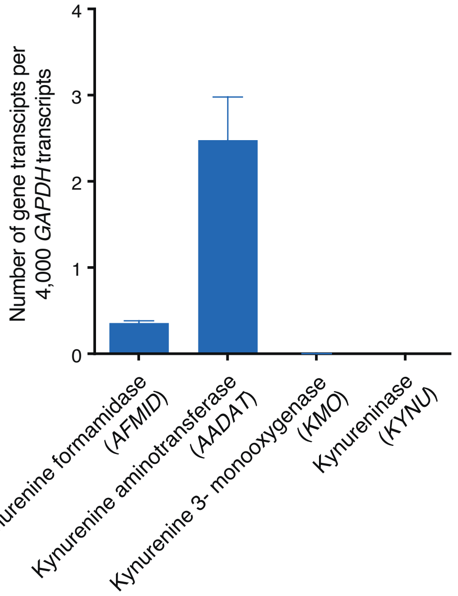


**Supplementary Figure 6: Tryptophan deprivation induces LAT1 expression via the GCN2 pathway**
**A.** Flow cytometric evaluation of kynurenine uptake in HEK293-E cells. HEK293-E cells were treated with different concentrations of kynurenine in PBS for 4 minutes at 37 C°. The reaction was stopped after 4 minutes by adding 125 μ l 4 % PFA for 30 minutes at room temperature in the dark. Data was acquired using 405 nm excitation (violet laser) and bandpass filter 525/50 BP (Bv510) on BD LSRII (Fortessa).
**B.** Western blot analysis of LAT1 expression for Fig **4G**.
**C.** qRT-PCR analysis of the expression of enzymes *AFMID (kynurenine formamidase), AADAT (kynurenine aminotransferase), KMO (kynurenine 3-monooxygenase), and KYNU (kynureninase)*, which are involved in kynurenine metabolism, in HEK293-E cells. Mean + SD of technical triplicates from one representative experiment out of three independent experiments. **** significant with p<0.0001, *** significant with p<0.001, ** p<0.01, * p<0.05 and NS= not significant.

Supplementary Figure 7

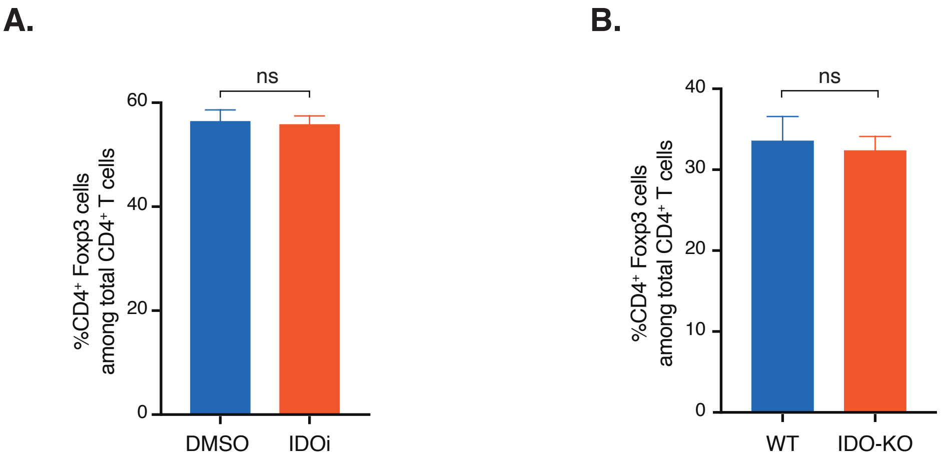


**Supplementary Figure 7: Impact of IDO on CD4+ FoxP3+ Treg-cell differentiation**
**A.** Treg differentiation (same as Fig 5D) of naïve CD4+ CD62L+ T cells isolated from WT mice treated or not with IDO1 inhibitor (Epacadostat, 2.5$\mu$M) during Treg differentiation. **B.** Treg differentiation (same as Fig 5D) of naïve CD4+ CD62L+ T cells isolated from WT or Ido1-KO mice. Mean + SD of technical triplicates from one representative experiment out of three independent experiments. **** significant with $p<0.0001$, *** significant with $p<0.001$, ** $p<0.01$, * $p<0.05$ and NS= not significant.
